# The cost of enforcing a marine protected area to achieve ecological targets for the recovery of fish biomass

**DOI:** 10.1101/216598

**Authors:** Christopher J Brown, Brett Parker, Gabby N Ahmadia, Rizya Ardiwijaya, Purwanto, Edward T Game

## Abstract

Protected areas are the primary management tool for conserving ecosystems, yet their intended outcomes may often be compromised by poaching. Consequently, many protected areas are ineffective ‘paper parks’ that contribute little towards conserving ecosystems. Poaching can be prevented through enforcement and engaging with community members so they support protected areas. It is not clear how much needs to be spent on enforcement and engagement to ensure they are frequent enough to be effective at conserving biodiversity. We develop models of enforcement against illegal fishing in marine protected areas. We apply the models to data on fishing rates and fish biomass from a marine protected area in Raja Ampat, Indonesia and explore how frequent enforcement patrols need to be to achieve targets for coral reef fish biomass. Achieving pristine levels of reef fish biomass required almost year-round enforcement of the protected area. Surveillance of the protected area may also be enhanced if local fishers who support the reserve report on poaching. The opportunity for local fishing boats to participate in surveillance was too small for it to have much benefit for total reef fish biomass, which increases slowly. However, specific functional groups of fish have much higher population growth rates and their biomass was predicted to increase markedly with community surveillance. We conclude that budgets for park management must balance the cost of conducting frequent patrols against supporting alternative activities, like education to build community support. Optimized budgets will be much more likely to achieve ecological targets for recovering fish biomasses and will contribute to fiscal sustainability of protected areas.

## Introduction

Protected areas are a primary tool for conserving ecosystems. Protected areas are often used to protect marine species from the effects of fishery exploitation, which reduce the biomass and diversity of species (Edgar et al. 2014). Recent international commitments to meeting Convention on Biodiversity targets has seen rapid growth in marine protected areas globally, with coverage increasing more than four times since 2000 (Watson et al. 2014, Boonzaier and Pauly 2016). However, many of these new protected area may be ‘paper parks’ that are not enforced (Gill et al. 2017). Globally, the marine protected areas with the highest biomasses and diversity of large fish are those that are old, large, fully protected from fishing, isolated and well enforced (Edgar et al. 2014).

Ensuring that protected areas deliver their intended conservation outcomes requires sufficient ongoing funding for enforcement and for building community support (Gill et al. 2017). The expense of enforcing protected areas may be a major impediment to their long-term success (Ban et al. 2011). Poaching in protected areas can erode their benefits for conserving biodiversity (Bergseth et al. 2015, Rizzari et al. 2015). Poaching may occur when poachers perceive the probability of detection is low and/or if the park’s objectives lack community support (Arias and Sutton 2013, Bergseth et al. 2017). Patrols of protected areas are critical to maintain compliance (Kelaher et al. 2015), but often budgets for patrols are not sufficiently resourced and patrols are not comprehensive enough to maintain compliance. Community support is also critical, so that fishers avoid poaching and report offenders. Community support can be achieved through engagement activities, such as education and consultation with communities on management plans (Leisher et al. 2012). However, the connection between expenditure on enforcement and the benefits of protection are generally not considered during the design stage, where the expectation around benefits typically involves an implicit assumption of perfect compliance (Davis et al. 2015). Numerous studies have addressed the opportunity costs of marine protected areas for fishing (e.g. Smith et al. 2010). What has not been addressed is how much needs to be spent on enforcing reserves so that fish biomasses are sufficient to conserve their ecological functions. Further, budgets for enforcement and community engagement are typically allocated ad-hoc, but budget allocations may be more effective if we could value community support in terms of avoided cost of patrols (Fox et al. 2017).

Here we develop an analytical framework for estimating the cost of enforcing protected areas so that fish biomass meets conservation targets. We estimate the cost of achieving specific biomass targets, including ecological relevant targets for fish biomass, where cost is given in general terms of days of patrols required. We apply the framework to model the Kofiau and Boo Islands Marine Protected Area in Raja Ampat Indonesia (Ahmadia et al. 2015). Raja Ampat is the global center of coral and fish diversity, but faces considerable pressure from fisheries. Efforts over the past ten years to establish protected areas have been successful and now management is transitioning to fiscal sustainability, thus quantifying budgetary needs for effective management is timely.

## Methods

First we describe a model of fish biomass inside protected areas, when the fish population is subject to variable levels of poaching. Then we describe application of the model to the case-study in Raja Ampat.

### Models of poaching, enforcement and compliance

We modeled poaching as a discrete and intermittent event, rather than using the traditional approach of modeling fishing mortality as a continuous pressure. Poaching events may often be intermittent, because poachers are fishing intensively for small amounts of time in an attempt to avoid enforcement officials. For instance, reefs in Indonesia are subject to fishing by ‘roving bandits’, commercial scale vessels that roam large areas and intensively fish local areas for relatively short-periods of time (often with illegal fishing gear), before they move to the next reef (Berkes et al. 2006). Small-scale poachers may also fish intermittently, for instance poaching by recreational fishers in the Great Barrier Reef marine protected area is most likely to occur on public holidays (Bergseth et al. 2017).

We developed two complementary models of poaching, which represent alternative plausible processes about the behavior of poachers. In both models we assumed fish growth was logistic growth with fixed parameters r (intrinsic growth rate) and *K*(maximal biomass), that poaching occurred at random intervals where the mean interval time *d* was described by an exponential distribution with rate *u*_*z*_ = 1/*d*. Thus, the equilibrium state for both of these models was a distribution of fish biomass.

The probability of a given biomass was calculated slightly differently for each model. In the first model, a poaching event ends once fish biomass has been depleted to a fixed level, *B*_0_ (Fig 1A). Fishing would deplete biomass to a fixed level if the marginal cost of harvesting fish increases as density in the reserve is depleted (e.g. White et al. 2008). Once costs exceed the expected revenue generated from poaching a reserve, a roving poacher will move elsewhere. This model had an analytical solution for the probability of different fish biomass levels ((Possingham 1989), Fig 1B). Taking the above assumptions for model one, we can calculate the probability of observing fish biomass *B*_*obs*_ at a random sampling time greater than or equal to a pre-specified level (*B*_*Q*_) as:

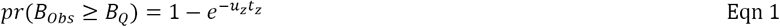

**Figure 1.**
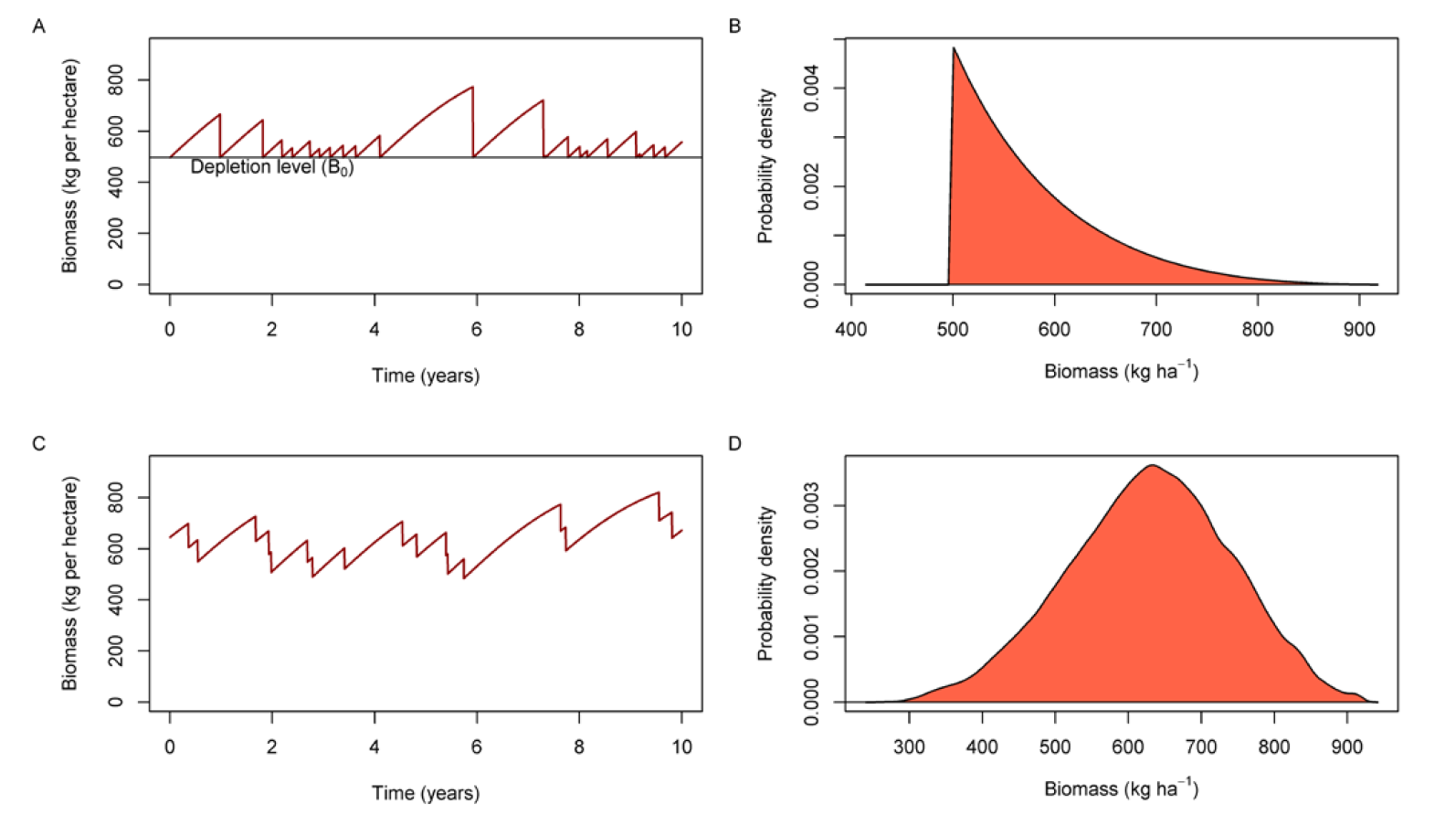
Example of fluctuations in fish biomass over time under model one (fixed depletion level A) and model two (fixed fishing rate C) and the probability density for biomass observed at random times for model 1 (B) and model 2 (D).

Where *t*_*z*_ is defined by the solution to the logistic growth function:

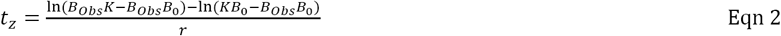

In the second model we assumed poaching mortality occurred at a fixed rate. This model was defined by the difference equation:

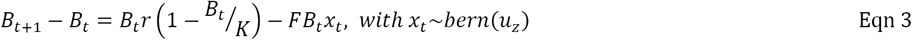

with *x*_*t*_ ∊ {0,1}

Where *F* is the fixed poaching rate and *x*_*t*_ is an indicator variable for whether poaching happened or not, and is drawn from a Bernoulli distribution with probability *u*_*z*_. Interval times between poaching events will follow an exponential distribution, as for model one. The mean poaching rate is *Fu*_*z*_ and in the limit when *u*_*z*_ =1 this model reverts to the difference form of the logistic model with a continuous harvest rate. Because the biomass after depletion in model two depended on when poaching started, we used simulations to determine the distribution of fish biomass. Simulations were run for 500 years on a daily time-step to ensure the distribution of fish biomass had converged on its equilibrium state.

Our next aim was to determine how enforcement affected the distribution of fish biomass. In both models, enforcement increased the average time interval between poaching events, such that enforcement patrols decreased the rate of poaching in proportion to the number of days per year that were patrolled:

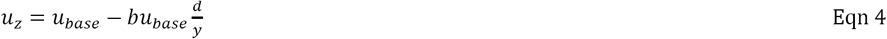

Where *u*_*base*_ was the poaching rate with no enforcement, *d* is the annual number of days that were patrolled, *y* is units per year (e.g. days = 365) and *b* is a parameter controlling how sensitive poachers are to enforcement. If *b* = 1 then poachers reduce their rate of poaching in proportion to the amount of enforcement. If *b* < 1 poachers are less sensitive to the rate of enforcement patrols. This may occur if poachers do not know of the park’s existence, are able to avoid detection, or penalties are insufficient (Byers and Noonburg 2007). If poachers are risk averse then b>1 and the rate of poaching decreases faster than the rate of enforcement.

Community support for a park may also increase the days patrolled, if community members engage in surveillance. Therefore, the final term in equation 4 for the proportion of days patrolled becomes:

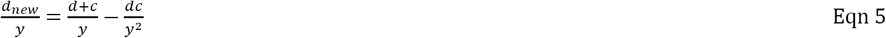

where *d* is the number of enforcement patrols per year and *c* is the number of days per year that community members are likely to report poachers if they encounter them. The model assumes community visits and patrols are independent and the term *dc*/*y*^2^ accounts for days when community visits and patrols co-occur.

### Application of the models to Raja Ampat marine protected areas

We applied the enforcement models to estimate the number of patrols per year required to achieve biomass targets for reef fish biomass in the Kofiau and Boo Island Marine Protected Area, Raja Ampat, Indonesia (fig. 2). Initially we presented results from a base-case, then we conducted further analyses to explore how the spatial and biological context of a reserve may affect the required rate of patrols. Our objective in these analyses was to explore the effect of different assumptions and contexts on the days patrolled, so we focus on comparing different scenarios and do not provide precise error estimates for days patrolled.

**Figure 2.**
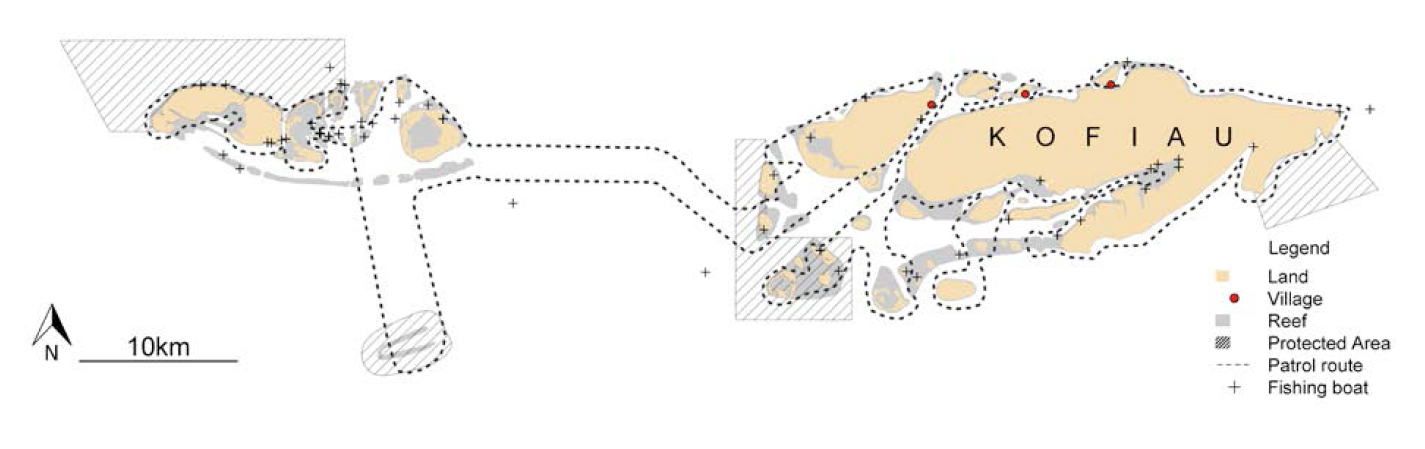
Map of the study region showing the patrol routes (which were assumed to be the same as the resource use monitoring survey routes), coral reef, protected areas and non-local fishing boats. Note that the protected areas were implemented after the resource use monitoring surveys.

In the base case we derived parameters for the models to represent reef fish biomass across all diurnally active fish observed in diver surveys, because total reef fish biomass is a common management target (MacNeil et al. 2015). We used underwater visual surveys of reef fish biomass to derive estimates of the depletion level (model one) and the fishing mortality rate (model two) (Table 1). Fish biomass 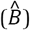 was estimated from diver surveys of reef fish biomass conducted at 39 sites around Kofiau’s coral reefs (Glew et al. 2015). To account for spatial variation in fish biomass across reefs, we interpolated mean observed biomass values per hectare across all reefs with a generalized additive model using a thin plate regression spline specified as an interaction over the x and y coordinate space (maximum degrees of freedom = 34) (Wood 2003). We could then estimate the regional fish biomass and mean biomass per hectare.

**Table 1.**
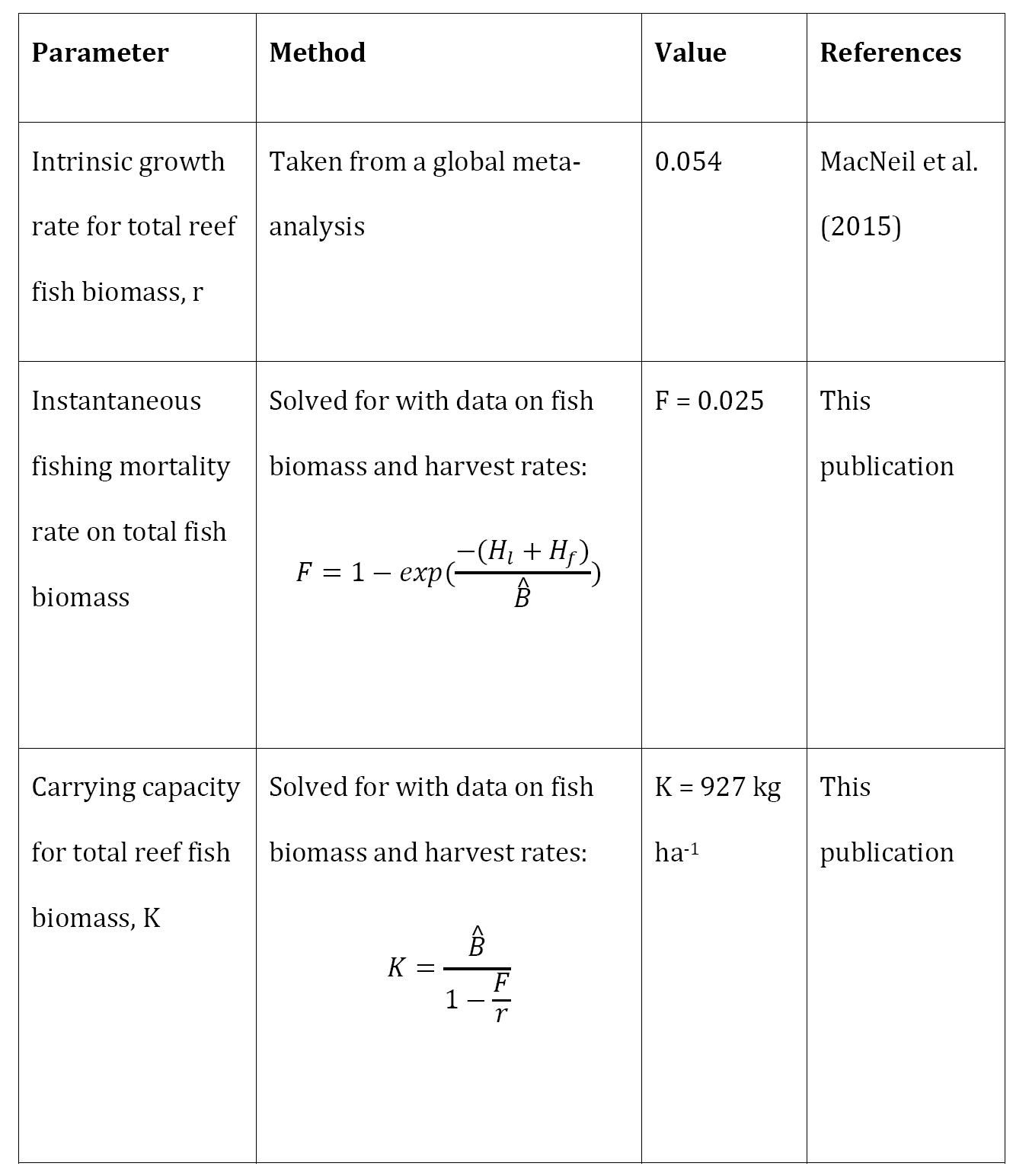
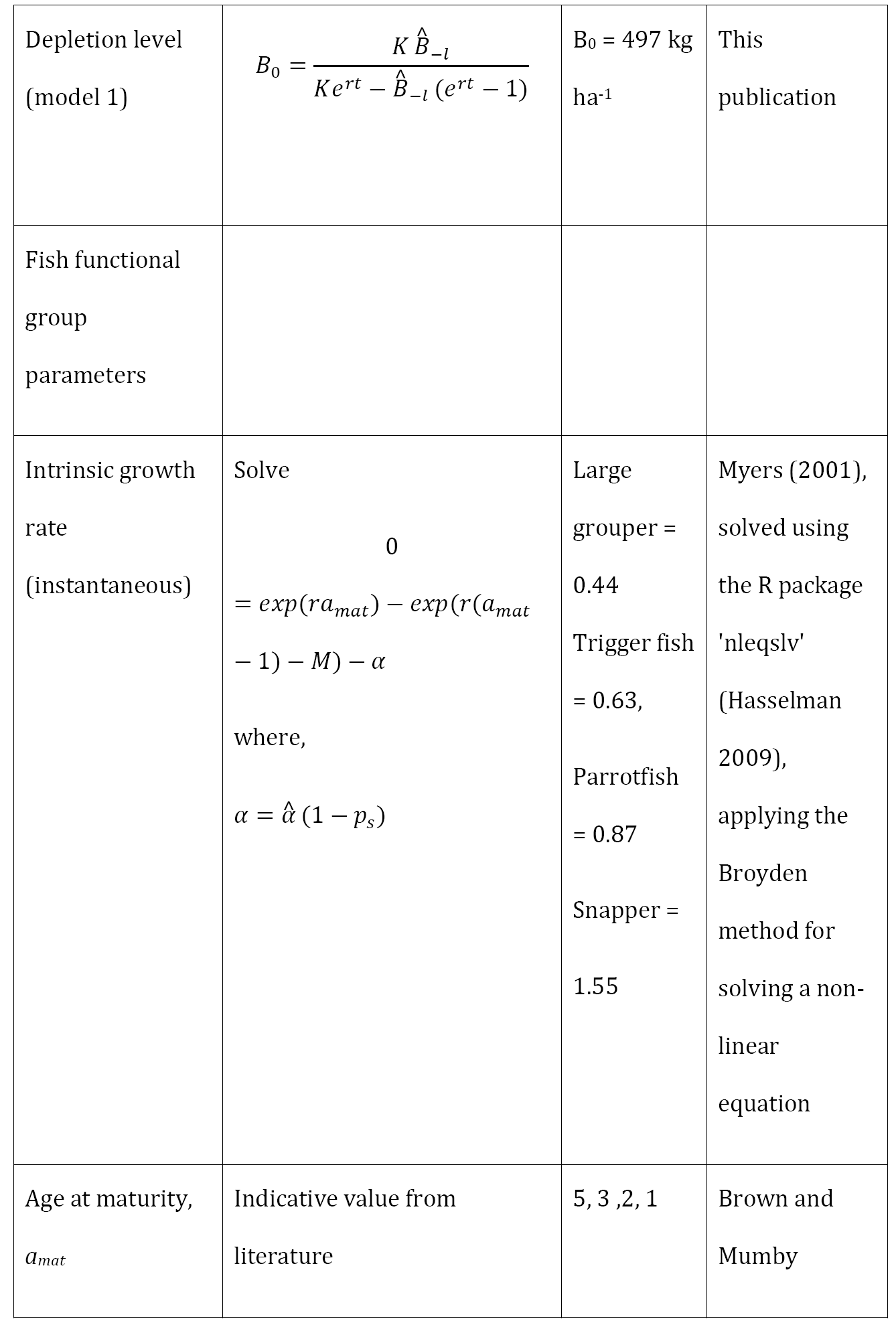
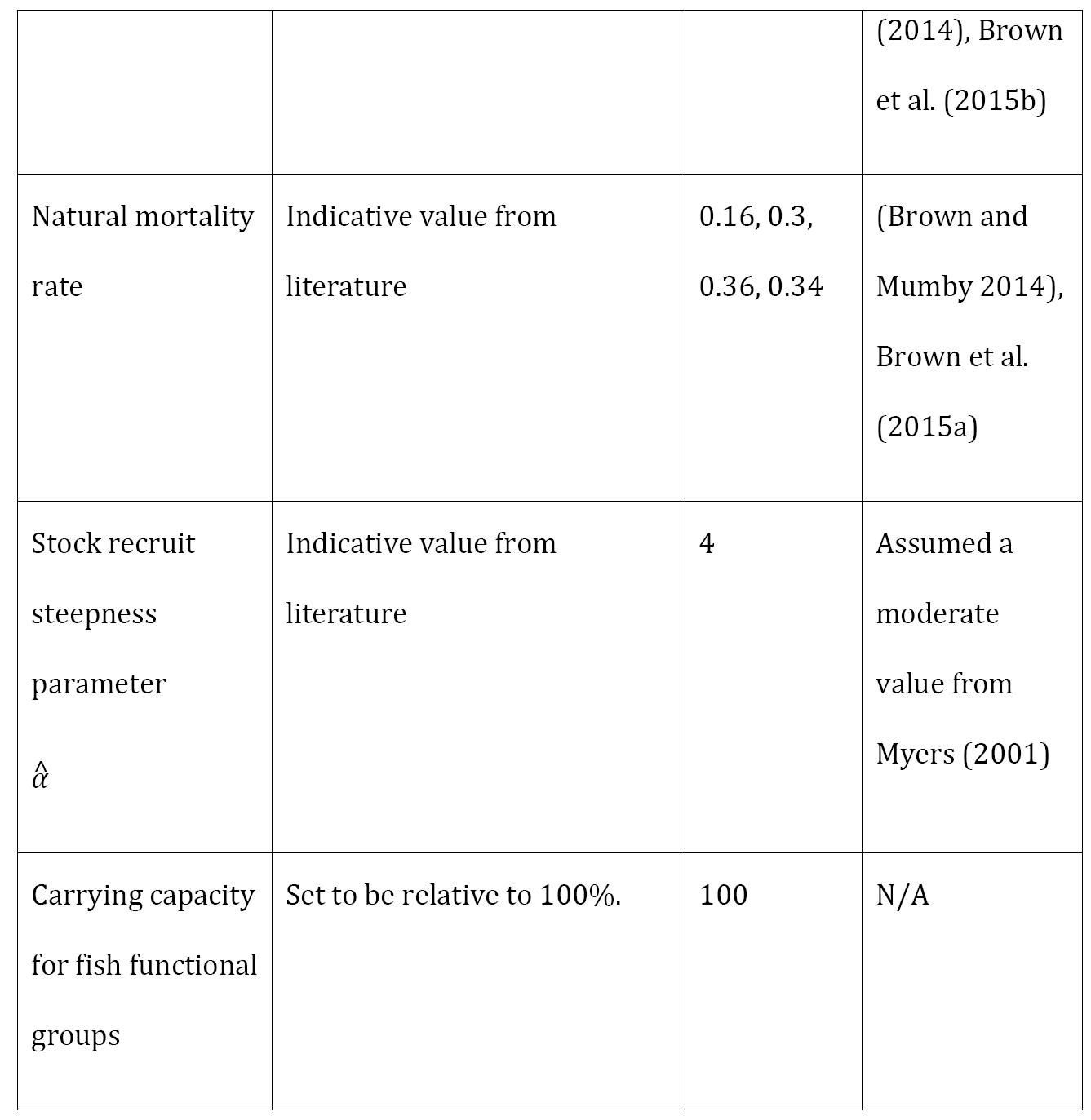
Methodology and references used to estimate model parameters. *H*_*l*_ and *H*_*f*_ were the annual harvest (per hectare) for local and non-local boats, 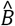 is the per hectare biomass observed on reefs 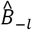 is the expected per hectare biomass without local fishing, *F* is the instantaneous fishing mortality rate and *p*_*s*_ is annual adult survival.

We used data from monitoring of resource use to estimate the baseline poaching rate. The resource use monitoring program conducted surveys around the island of Kofiau from 2005 to 2009 (36 surveys), before the protected areas were officially decreed. During surveys a patrol boat drove a predetermined route and approached all boats that were observed (fig. 2). Each boat was surveyed, including type of boat, type of fishing gear, activity (travelling, resting or actively fishing), catch if any and home port. It was assumed that since the implementation of the MPAs fishing rates inside the MPAs would be equal to or less than this baseline. We only analysed poaching by non-local boats (those from outside of the Papua region). Poaching by locals is likely negligible because they generally support the MPAs, for instance they are involved with monitoring parks and report on illegal fishing (Fox et al. 2017). Local catch is also relatively small, making up 22% of the legal catch with a mean catch per local boat of 3.4 kg (79% of catches were <5kg). Non-local boats are typically larger and will visit region for a limited period of time and take a large catch, for instance their mean catch was 231 kg and 16% of boats surveyed had caught > 500kg. Many of these non-local boats may operate like roving bandits, moving across large spatial regions and sequentially depleting coral reefs as they move.

In the second set of analyses we estimated the value of surveillance by local fishers to increasing fish biomass in reserves. We therefore needed to estimate the number of local boats in the vicinity of reserves whom might report on poaching. We expected the number of local fishing boats to vary with distance to villages and we estimated the potential number of days when local fishers were near MPAs at two distances from the villages, the maximum value near to villages (100 m) and the expected value far from villages (15 km). These values were chosen to bound the range of plausible values. We used the data for local fishing boats fishing on reefs from the resource use monitoring surveys. The number of boats within 600m x 600m grids at different distances from villages was estimated using a generalized additive model with a thin plate spline on distance to village (with a maximum degrees of freedom of 5). In total we had 105 observations of local fishing boats across 306 grid cells that ranged in distances from villages from 0.2 to 49.6km. Counts of boats in the grid cells were standarised by the number of days surveyed, then scaled up to annual estimates. A semivariogram of the residuals did not indicate any spatial autocorrelation once distance to village was included in the model, so we fitted the model without any autocorrelation structure. The model assumed Poisson errors for the response (number of boats per year). Then, to estimate the potential number of days that community members would incidentally report on poaching, we used the model to estimate the annual visitation rate by local boats for a protected area that had a size of size of 20 hectares, the typical size of no-take protected areas in the region.

In the third set of analyses, we explored the effects of changing the sensitivity of poaching to enforcement. The sensitivity parameter is unknown, and would be challenging to estimate, so here we present results for three values of *b* that cover a range of plausible values. The first was b=1, which assumes that poaching rate declines in proportion to the days patrolled. We compared this scenario to two others where *b* was increased by 50% (greater sensitivity) or reduced by 25% (lower sensitivity).

In the final set of analyses, we explored the sensitivity of key results to fish life-history types. We modelled four fish functional groups that cover a range of life-history parameters (Abesamis et al. 2014), have previously been identified to perform important ecological functions in coral reef ecosystems (Brown and Mumby 2014) and include some groups that are currently used as indicators for the park’s status (Glew et al. 2015). The reef fish groups were: large groupers (family Serranidae) that predate on meso-predator fish and thus can suppress trophic cascades but are relatively slow growing; triggerfish (family Balistidae) that are important predators of bio-eroding sea urchins and have a moderate population growth rate; and parrotfish (family Scaridae) whose grazing can suppress algal blooms and are relatively fast growing (Brown and Mumby 2014). We also include snappers (family Lutjanidae) because they are an important indicator species for the Kofiau reserve (Glew et al. 2015). It was not possible to estimate the model parameters for the biomass and fishing rate of reef fish functional groups, because the resource use monitoring survey did not resolve fish to the Family level (Table 1). Therefore, we conduct sensitivity analyses to explore the effect of fishing rate and depletion level on enforcement costs.

## Results

For Kofiau, we estimated a pristine fish biomass of 927 kg ha-1. Total fish biomass was largely insensitive to increasing the days patrolled. Under model one fish biomass only increased when there were >340 patrol days per year. Slow growth also meant that many days of enforcement (>330) were required to see any noticeable increase in fish biomass for model one (fig. 3A).

**Figure 3.**
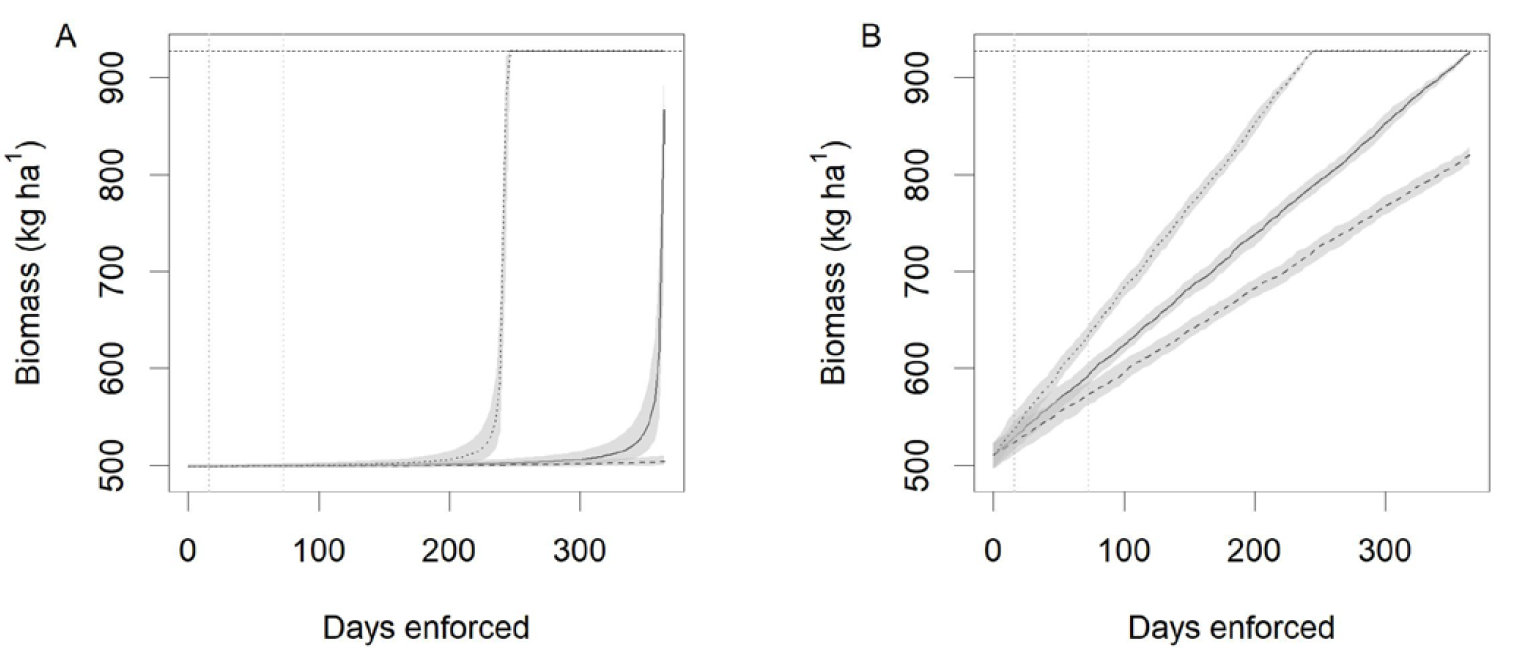
Fish biomass vs days patrolled for (A) model one (constant depletion level) and (B) model two (constant rate of fishing). Solid lines give b=1, dashed lines b = 0.75 and dotted lines b = 1.5. Horizontal dashed lines indicates the carrying capacity, vertical dotted lines indicate the number of equivalent enforcement days if community members engage in surveillance.

Under model two, fish biomass increased linearly with number of days patrolled. Reef fish biomass has a slow recovery rate (MacNeil et al. 2015), so we expected total fish biomass to be insensitive to the number of days patrolled when existing fishing rates are moderate. For instance, 340 days were required to attain a fish biomass that was 80% of carrying capacity under model one, or 203 days under model two.

The model of boats against distance to village indicated a significant decline in the number of boats further from villages (F = 371, p<0.001, effective degrees of freedom= 3.98, fig 4). Most fishing was observed within 20km of villages (50% of fishing boats were observed < 4.2 km from villages and 75% of boats were <17.5 km), though boats occurred as far away as 50km from villages (Fig 4). The predicted number of local boat days in a 20 hectare protected area per year was 13.1 (12.6-13.6 S.E.) at 1km from villages and 0.55 (0.53 – 0.57 S.E.) 15 km from villages. The maximum was 73 boat days per year.

**Figure 4.**
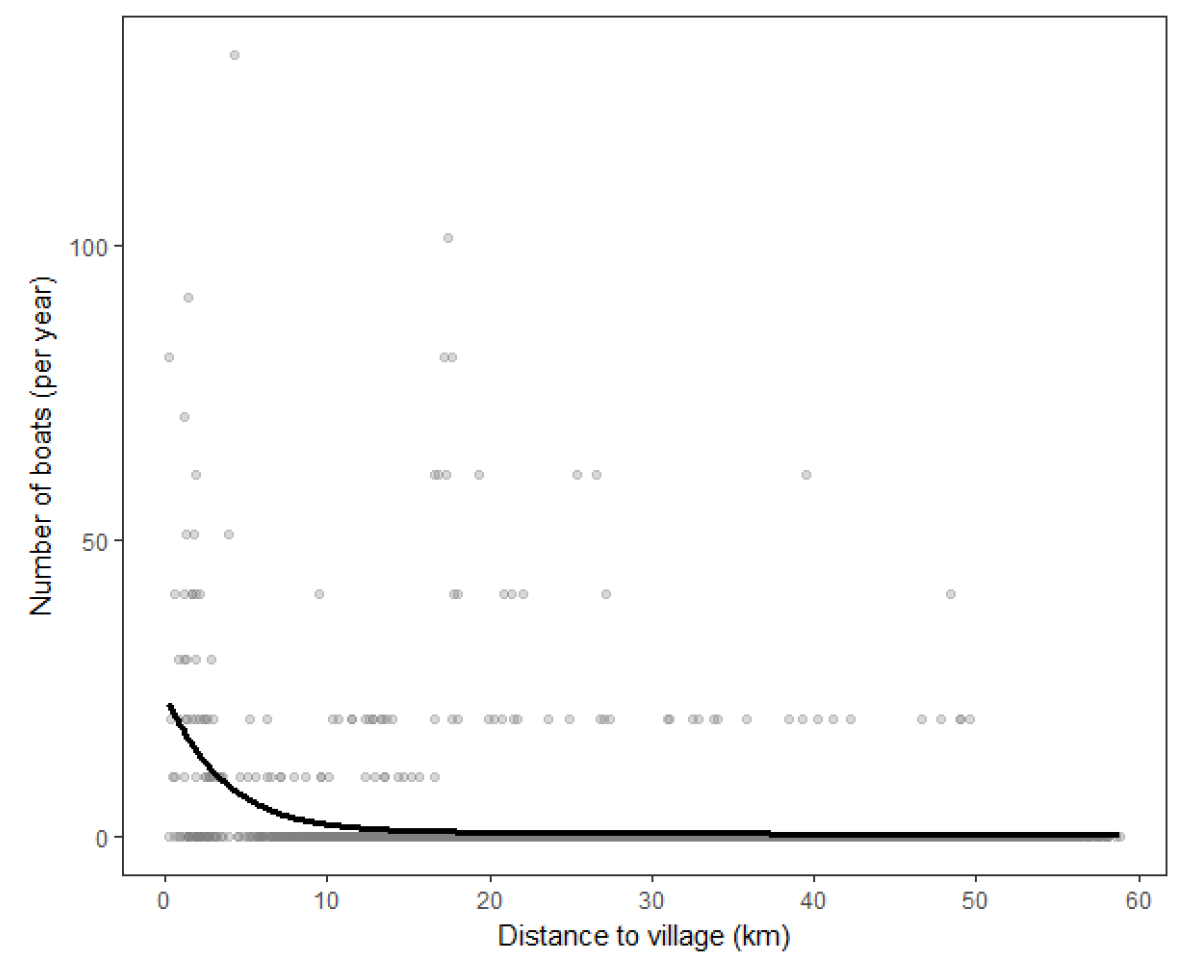
Effect of distance to villages on the number of local boat days of active fishing. Points show observations from surveys (standardized to be per year) and the black line shows the fitted mean from the generalized additive model.

The insensitivity of total fish biomass to enforcement meant community surveillance had only a small effect on fish biomass. For instance, even assuming all boats near to villages reported on poaching (73 boats per year), fish biomass was not predicted to increase under model one, and increased only from a mean of 510 kg ha^-1^ to 594 kg ha^-1^ under model two (fig. 3). At a distance of 15 km from a village, the potential benefits of community surveillance for fish biomass were small for total fish biomass.

Fish biomass was much greater if poaching had greater sensitivity to enforcement than assumed in the base case (Fig 3). For instance, increasing *b* by 50% reduced the number of patrol days required to achieve carrying capacity from nearly 365 to 246 under model one and from 365 to 239 under model two. If the sensitivity was reduced then it was impossible to achieve a carrying capacity biomass under either model (Fig 3).

The four fish functional groups had much higher population growth rates than total fish biomass (Table 1), so recovery of their biomasses was more responsive to increasing enforcement. The faster growth of the functional groups than total fish biomass meant that community participation in surveillance in combination with some patrols may enhance the biomass of reef fish functional groups. For instance, even with no enforcement, biomass was expected to be >50% of carrying capacity for all functional groups with the baseline level of fishing pressure (figs 5E and 6A), with greater value for faster growing functional groups. For model one, small incremental increases in biomass required significantly more enforcement (fig 5), because the fixed depletion level meant infrequent poaching could push biomass back to the baseline level. For model two, there was a linear increase in the number of days required to achieve higher biomass targets. The faster growth of the fish functional groups than total fish biomass meant that community participation in surveillance was expected to significantly enhance their biomass.

**Figure 5.**
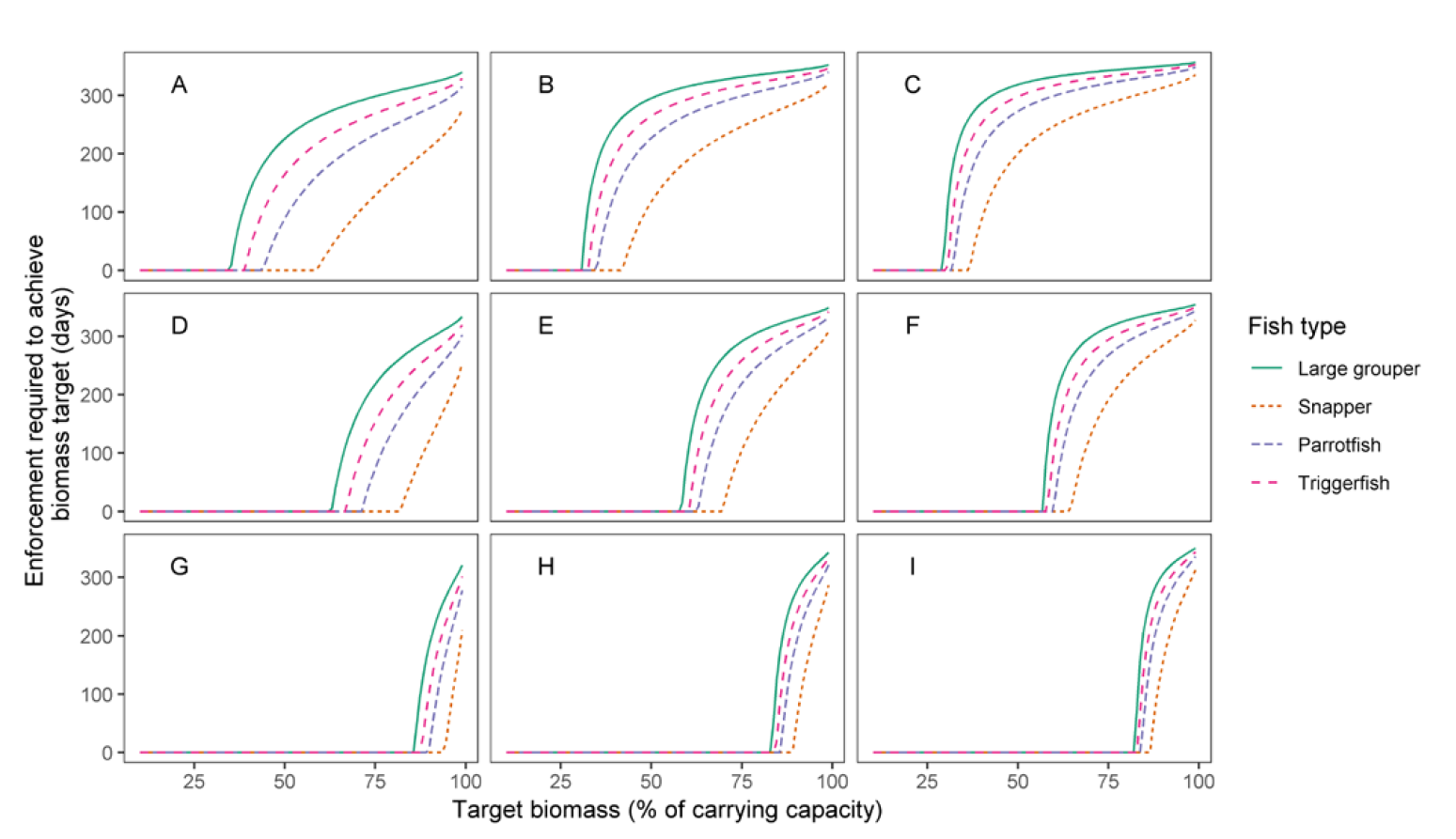
Days of enforcement per year required for achieving biomass targets for each fish functional group for: high (A, B, C), moderate (D, E, F) and little (G, H, I) depletion; and for low (A, D, G), moderate (B, E, H) and high (C, F, I) intervals between poaching events. (A) has the overall highest fishing pressure and (I) has the lowest. For each parameter the lower values are half the baseline, moderate values are equal to the baseline and high values are double the baseline, where the baseline are those values that were estimated for the primary analysis.

Of the four fish functional groups, the days of enforcement was greatest to achieve biomass levels relative to unfished for slow growing grouper and lowest for the fast growing snapper (figs 5 and 6). For all functional groups under model one, it required less enforcement to achieve the same biomass level when the minimum depletion level was higher (Fig 5 - compare rows) or the average interval between poaching events was greater (Fig 5 - compare columns). For instance, achieving biomasses of 50% of their unfished levels required 200 to 300 days of enforcement if the fish groups were depleted to low levels and there was frequent poaching (Fig 5C), whereas no enforcement was necessary to achieve fish biomasses >75% of their unfished level if poaching was infrequent and there was little depletion of fish biomass (Fig 5G). Similarly for model two, if fishing mortality rate (Fig 6) or the rate of poaching was increased, then a greater number of patrol days were required to achieve a given fish biomass target.

**Figure 6.**
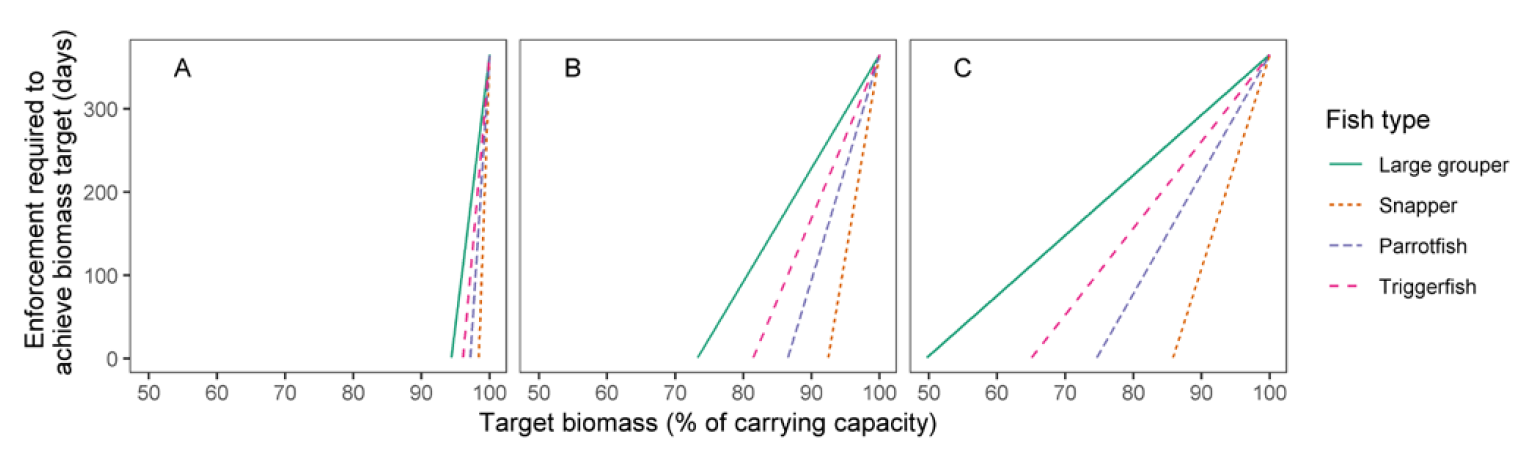
Days of enforcement per year required to achieve biomass targets for each fish functional group for low (A), moderate (B, 5 times baseline from Kofiau) and high (C, 10 times baseline for Kofiau) fishing mortality rates. Results for increasing the frequency of poaching events were similar to increasing the fishing mortality as shown here.

## Discussion

Estimating the cost of enforcing protected areas can help inform appropriate allocation of management resources and increase the likelihood of protection achieving its intended benefits (Ban et al. 2011, Davis et al. 2015). We estimated the number of patrol days required to achieve biomass targets for reef fish in an Indonesian reserve system under two models. Under both models, total reef fish biomass was relatively insensitive to an increasing rate of patrol, meaning a high frequency of patrols was required to achieve pristine fish biomass. Specific functional groups had faster population growth rates and their biomasses benefited more from enforcement when fishing pressure was high. The constant fishing rate model had higher biomass for an intermediate number of patrols than the fixed depletion level model. Fixed rate poaching would occur if fishers spend a similar amount of time during each poaching event, whereas fixed depletion would occur if poachers fish until fish density is too low for further fishing to be profitable. Therefore, the benefits of partial enforcement for fish biomass depend on the drivers of poaching pressure. Determining these drivers will be important to identify the cost of achieving a certain biomass target.

Gaining community support for protected areas may enable functional levels of fish biomass to be achieved with fewer patrols and at a much lower cost, because community members are less likely to poach themselves and may also participate in surveillance (e.g. Fox et al. 2017). In most cases effective protected areas will require both enforcement and community engagement (Watson et al. 2015, Bergseth et al. 2017). We found there were relatively few local boats operating near the protected areas in Kofiau, even if they were close to villages, so the effect of local surveillance was not predicted to bring much benefit to increasing total fish biomass. One caveat is that we did not account for shoreline observation, which could be almost year round if a protected area is located near a village or tourist resort. Accounting for surveillance by shoreline observation when designing a reserve network may significantly affect its final design. For instance, greater emphasis may be put on the importance of protected areas that are close to community centers, but future work should address how such benefits can be reconciled against lost opportunities for fishing by local community members (e.g. Davis et al. 2015).

The sensitivity of poaching to enforcement rate had a much greater effect on expected fish biomass than community participation in surveillance. Efforts to increase this sensitivity could offer a very cost-effective way to improve the effectiveness of enforcement. Engagement activities can improve compliance by increasing the perceived probability of detection of poachers by enforcement (Bergseth et al. 2017). Increasing the amount and rate of fines is also expected to enhance compliance among fishers who are well aware of the risk of being fined (Byers and Noonburg 2007, Kelaher et al. 2015). We did not consider the level of fines here, because we were modelling poaching by transient non-local boats, many of whom may not know about the existence of protected areas, so fines are unlikely to be an important driver of their behavior. There have been extensive efforts to increase awareness about Kofiau’s protected areas among local communities (Leisher et al. 2012), members of whom are now also employed to monitor and patrol protected areas (Fox et al. 2017). This model of engagement has resulted in much greater compliance from local people and may also help to deter roving bandits because local people who are invested in protecting the protected areas will aid in surveillance and enforcement of the protected areas when they are out fishing (Berkes 2010, Fox et al. 2017). Efforts to increase awareness about protected areas among non-local fishers may also be necessary to achieve full protection of the fish stocks.

Further work is needed to quantify how poaching rates are affected by community engagement activities, so that management budgets can be balanced between these activities. A total of $USD1,020,223 was spent over 2004-2010 leading up to and during the creation of the protected areas around Kofiau and around the nearby island of Misool (Leisher et al. 2012). This engagement considerably increased community understanding and support for the protected areas between 2004 and 2010 (Leisher et al. 2012). Awareness is important, because local fishers may poach unintentionally if they do not know of the reserve. This increase in community support may also reduce poaching by roving bandits through local people reporting illegal fishing to enforcement officers (Alder 1996). The key uncertainty in these estimates for the value of community engagement was the sensitivity of poaching rate to enforcement and how this changes when there is community supported for the protected area. Greater awareness of the protected area, and fines for non-compliance should increase the sensitivity of poaching to enforcement (Byers and Noonburg 2007, Kelaher et al. 2015). Social surveys that quantify how reporting and poaching rates change across users whose participation in engagement programs varies are now needed to more accurately quantify the value of engagement activities. Social surveys are thus an important part of programs that evaluate the impact of conservation actions (e.g. Glew et al. 2015, Fox et al. 2017). With these values in hand, our models could be used to value community support for park and thus apportion conservation budgets between engagement activities that enhance community support and enforcement.

The overall cost required to be spent on enforcement may be reduced if the target of patrols is to achieve biomasses of fish functional groups that are sufficient for them to perform ecological functions, rather than trying to achieve pristine levels of total fish biomass. Total reef fish biomass has a much slower growth rate than specific functional groups that are important for the Kofiau protected area, like snapper and parrotfish (Glew et al. 2015). Total reef fish biomass may increase slowly because of changes in the composition of fish communities during recovery (MacNeil et al. 2015). However, achieving functional levels of biomass may in some cases require significantly less enforcement than attempts to achieve pristine fish biomass. For instance, triggerfish may need to be at 80% of their unfished biomass to provide their functional role of predating on bioeroding invertebrates (Brown and Mumby 2014). We estimated that it may require ∼150 days of enforcement to achieve an 80% triggerfish biomass if fishing pressure on this group is similar to the ecosystem aggregate estimate of fishing pressure.

Our model can be used to help inform budgeting for management of protected areas, both to estimate how much is required to support a park and also how to allocate a budget between community engagement versus enforcement. Future studies should combine dynamic estimates of sufficiency like ours with more complex spatial algorithms for developing MPAs (Davis et al. 2015) and more complete estimates of enforcement cost (Ban et al. 2011). Considering the cost of effective protected areas is important in the planning process, for instance, estimates of the cost can be used to focus development of new protected areas in places where they will be most effective. Ultimately, both community support and enforcement are necessary to effectively conserve the biodiversity in protected areas.

## Acknowledgements

CJB was supported by a Discovery Early Career Researcher Award (DE160101207) from the Australian Research Council. We are grateful to four reviewers for helpful comments.

